# Machine Learning for Prioritization of Thermostabilizing Mutations for G-protein Coupled Receptors

**DOI:** 10.1101/715375

**Authors:** S. Muk, S. Ghosh, S. Achuthan, X. Chen, X. Yao, M. Sandhu, M. C. Griffor, K. F. Fennell, Y. Che, V. Shanmugasundaram, X. Qiu, C. G. Tate, N. Vaidehi

## Abstract

Although the three-dimensional structures of G-protein-coupled receptors (GPCRs), the largest superfamily of drug targets, have enabled structure-based drug design, there are no structures available for 87% of GPCRs. This is due to the stiff challenge in purifying the inherently flexible GPCRs. Identifying thermostabilized mutant GPCRs via systematic alanine scanning mutations has been a successful strategy in stabilizing GPCRs, but it remains a daunting task for each GPCR. We developed a computational method that combines sequence, structure and dynamics based molecular properties of GPCRs that recapitulate GPCR stability, with four different machine learning methods to predict thermostable mutations ahead of experiments. This method has been trained on thermostability data for 1231 mutants, the largest publicly available dataset. A blind prediction for thermostable mutations of the Complement factor C5a Receptor retrieved 36% of the thermostable mutants in the top 50 prioritized mutants compared to 3% in the first 50 attempts using systematic alanine scanning.

**Statement Of Signifigance:** G-protein-coupled receptors (GPCRs), the largest superfamily of membrane proteins play a vital role in cellular physiology and are targets to blockbuster drugs. Hence it is imperative to solve the three dimensional structures of GPCRs in various conformational states with different types of ligands bound. To reduce the experimental burden in identifying thermostable GPCR mutants, we report a computational framework using machine learning algorithms trained on thermostability data for 1231 mutants and features calculated from analysis of GPCR sequences, structure and dynamics to predict thermostable mutations ahead of experiments. This work represents a significant advancement in the development, validation and testing of a computational framework that can be extended to other class A GPCRs and helical membrane proteins.

## Introduction

GPCRs, the largest superfamily of drug targets, reside in membranes, where they coordinate agonist stimulation with activation of different G-proteins and/or β-arrestin signal transduction pathways. Apo and ligand-bound GPCRs exhibit a dynamic equilibrium among multiple functional conformational states(Thal et al., 2018) broadly classified as active and inactive states. The active state conformations of class A GPCRs are characterized by the large movement of the intracellular regions of transmembrane (TM) helices TM6, TM5 and TM7 as illustrated in Fig. 1A using the crystal structures of adenosine receptor A_2A_R. The relative population of these conformation states are modulated by binding of ligands and G-proteins or other intracellular transducers(Bhattacharya and Vaidehi, 2010; Kim et al., 2013; Niesen et al., 2011). Therefore, to enable structure-based ligand design for GPCRs, it is imperative to solve the three-dimensional structures of GPCR conformations with various ligands and/or intracellular transducer proteins bound. However, the conformational flexibility of GPCRs poses significant challenges in stabilizing and purifying these proteins for structural studies. The breakthrough in protein engineering technologies in the past decade has resulted in a bounty of GPCR structures in inactive and active states that have provided many new functional insights(Tautermann and Gloriam, 2016).

**Figure 1:**
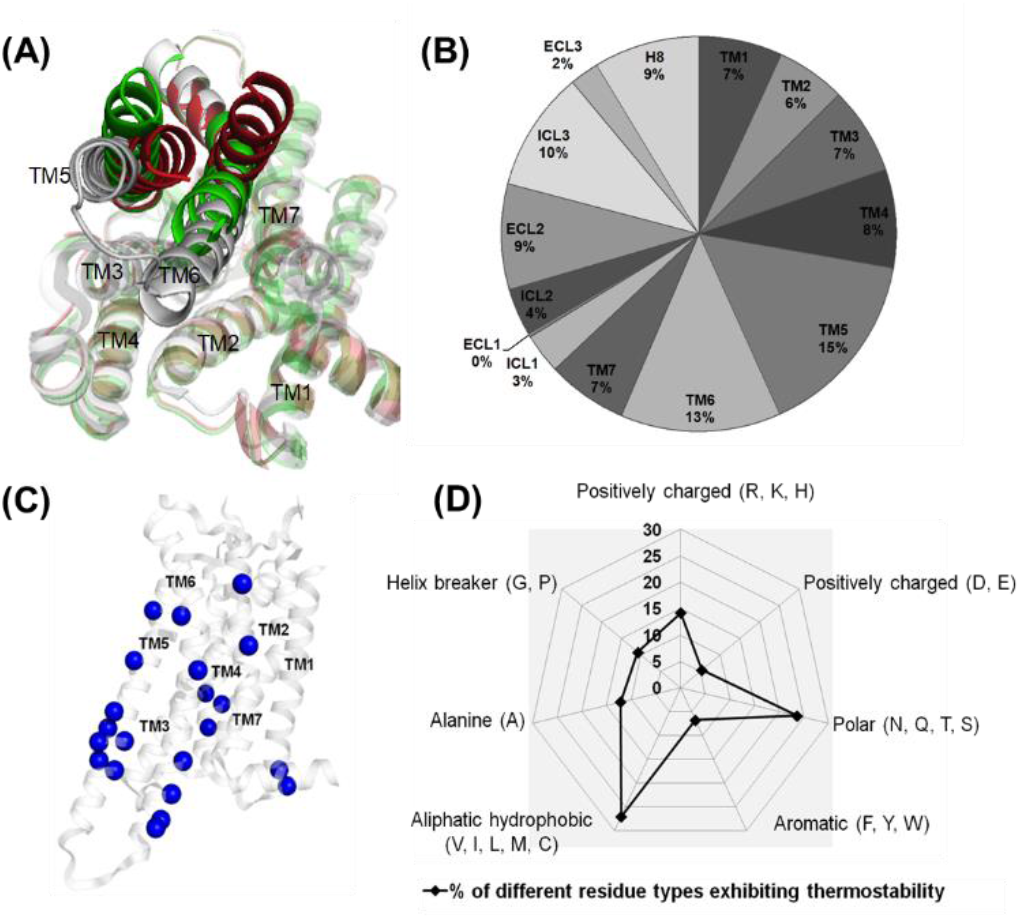
Analysis of the experimental thermostability data on 1231 thermostable GPCR mutants. **A**. The overlay of the crystal structures of A_2A_R in the antagonist bound inactive (grey), agonist bound active-intermediate (green) and agonist and G-protein bound fully active state (red). The intracellular view shows large movements in TM5, TM6 and TM7 upon activation. **B.** Analysis of the location of the thermostable mutant positions in all the five experimental thermostability datasets. **C.** Location of the thermostable mutants in the transmembrane regions that are common in β_1_AR, A_2A_R inactive and active-intermediate states and in NTSR1 active-intermediate states. **D**. Classification of the thermostable mutants by the amino acid properties.

One such protein engineering technology that yielded multiple GPCR structures is via thermostabilization, pioneered by Tate and coworkers(Tate and Schertler, 2009). This method involves performing systematic alanine scanning mutations to identify thermostabilizing mutant positions in the GPCR. These positions can be subsequently combined to yield a thermostable mutant that exhibits conformational homogeneity, enabling purification of a selected GPCR conformational state. Although this strategy has been successful for purifying multiple GPCRs in various conformational states(Lebon et al., 2011; Magnani et al., 2008; Serrano-Vega et al., 2008; Shibata et al., 2009), deriving an optimal combination of thermostable mutations quickly escalates cost and scale of experiments/research and development. Therefore, this procedure is only accessible to a handful of laboratories worldwide and requires smarter methods to democratize this process. Computational methods that can reliably predict the thermostabilizing mutations of a GPCR ahead of experiments greatly reduce the burden by prioritizing only promising alanine mutations. Our prior studies on properties of thermostable mutant GPCRs showed that energy function calculated using all-atom forcefields and a conformational ensemble rather than a single structural model, recapitulates thermostability and recovers more thermostabilizing mutations in the predictions(Bhattacharya et al., 2014).

Building on this success, in this work we trained four different machine learning (ML) models on the largest publicly available thermostability data for 1231 GPCR mutants from four different class A GPCRs. The ML models were trained using a combination of 26 features calculated from (a) amino acid sequence covariation analysis of all class A GPCRs, (b) GPCR structural features extracted using network analysis, and (c) thermodynamic energies calculated from an ensemble of conformations of the GPCRs that include a membrane potential. We performed a blind prediction of thermostabilizing mutations for the human Complement C5a Receptor, C5aR. We recovered 14 out of 39 thermostable mutants within the top 50 prioritized mutations, while systematic alanine scanning mutation has a probability of only capturing 1 mutant in 50 attempts. 4 out of the 14 thermostable mutants showed strong thermostability. We present a robust machine learning workflow that is extensible to other class A GPCRs. This makes the predictions of thermostabilizing mutations for any class A GPCR accessible to the larger community.

## Methods

### Experimental Thermostability Measurements

To train the machine learning models, we have used the thermostability data on 1231 single alanine scanning mutants on four class A GPCRs (two sets of data for inactive state and active-intermediate state stabilization for A_2A_R) measured by Tate and coworkers. We have thermostability data only for 200-250 mutations in each of the four GPCRs tested. For training the machine learning algorithms, we have collected features from our computational models that correspond to these residues for which thermostability data is available in the training dataset. The experimental measurements were performed on antagonist cyanopindolol bound turkey β_1_-adrenergic receptor in the inactive state (Serrano-Vega et al., 2008), agonist 5’-*N*-ethylcarboxamidoadenosine (NECA) bound active-intermediate state of adenosine receptor A_2A_R (Lebon et al., 2011) and antagonist ZM-241385 bound inactive state of A_2A_R (Magnani et al., 2008), agonist NTS1 bound rat neurotensin receptor NTSR1 bound (Shibata et al., 2013) and antagonist ZD7155 bound human angiotensin receptor 1, AT1R Briefly, the experimental thermostability was measured by heating the detergent-solubilized mutant to an elevated temperature (~28-32°C), cooling to 0°C, and the amount of correctly folded receptor was determined by a radioligand (either an agonist or antagonist or both) binding assay. In the work presented here, we have defined the thermostability of the wild type receptor as 100% and any mutants that showed 100% or higher ligand binding compared to the wild type receptor are defined as thermostable. For the β_1_AR, A_2A_R and AT1R receptor, the mutants were heated without any ligand present. In the case of NTSR1, two types of experiments were performed, one where the mutants were heated in the presence of the agonist neurotensin, and the other experiment heated the receptor in the absence of neurotensin, the data that is used in our machine learning models only included data from heating without the presence of agonist.

### Thermostability measurements for C5aR

The wild-type human C5aR receptor gene was cloned into mammalian cell expression vector pcDNA3.1 with the haemagglutinin (HA) signal sequence followed by a Flag epitope tag at the N terminus and a 10× His-tag at the C terminus. A library of 283 mutant C5a receptors was constructed with individual residues from Val35 to Leu311 of C5aR mutated to alanine or leucine (if the original residue is an alanine) by PCR-based site-directed mutagenesis. Each clone of the mutation was transient transfected in 6ml of human HEK-293 Expi cells and 6 million cells harvested after 2 days expression were pelleted in aliquots and frozen for thermostability screening. Six million cells expressing wild type-C5aR or mutant C5aR were suspended into 650 ul of PBS supplemented with protease inhibitor cocktail. Thermostability of C5aR mutants was assessed in a high-throughput three-temperature assay screening and all experiments were run in triplicate. For non-heat-treated samples aliquots of cell suspension were kept on ice while other aliquots of cell suspension were heated at 43 C (apparent Tm determined previously) and 46°C for 15 min respectively and the samples were then cooled on ice for 5 min. The following radioligand binding assay was done in a 96-well format; the functional expression level of each C5aR mutant was assessed on non-heat-treated samples and the thermal stability on thermal treated samples in their cellular lipid environment. In each 200 μl of radioligand binding assay reaction, 20 μl of C5aR cellular suspension was incubated in binding buffer (75 mM Tris-HCl pH 7.4, 1 mM EDTA, 5 mM MgCl2, 100 mM NaCl) with 100 nM [^3^H] Compound 1 (PF-06733901) at room temperature for 1 hour. Nonspecific binding of [^3^H] PF-06733901 was determined in the presence of 2.5 μM unlabeled ligand. Unbound radioligand was then removed from the reaction on PerkinElmer Filtermate and radioactivity was counted by liquid scintillation on a PerkinElmer Microbeta. Data were analyzed using Prism software (GraphPad).

### Partitioning the experimental data into training and testing ensembles

To train the machine learning models, we have used the thermostability measured by Tate and coworkers on every residue in four different class A GPCRs listed above. Experimentally we classify a mutant as thermostable if the experimental thermostability value is at or above the wild type (WT) threshold which is a score of 100. To convert this experimental data into labels for training machine learning models we used threshold ranking. If a mutation has a measured thermostability score of 100 (the score for the wild type receptor) or more it is considered thermostable and given a label “1”. Any score below 100 is considered non-thermostable and given a label “0”. These labels are used to classify our computationally calculated dataset.

### Balancing the training data

The experimental data is heavily imbalanced towards the non-thermostable mutants. In most cases the number of non-thermostable mutants is 3 times more than thermostable mutations. The 85% of the thermostability data that we used for training was balanced using a method called the Synthetic Minority Oversampling Technique (SMOTE) SMOTE-Tomek sampling. With SMOTE-Tomek method we were able to both under sample the majority class (non-thermostable mutants) and oversample the minority class (thermostable mutants) simultaneously. SMOTE oversamples the minority class using its k nearest neighbors (KNN). Tomek links removes samples from the majority class by removing outliers between the two classes using distance measurements. SMOTE forms new minority class(thermostable mutants) data by interpolating between several minority class data points that lie together(Batista et al., 2004). The results of the effect of balancing the data are shown in sections below.

### Generating Features for the Machine Learning Algorithms

The quality of predictions made using machine learning algorithms requires accurate description of “features” that describe the property of the outcome (in this case thermostability of GPCR mutants). We have computed several physical properties of GPCR structures listed in Table S2 that contribute to its stability as features for training the machine learning algorithms (listed in Table S2).

### Energy Related Features

In our previous studies of GPCR thermostability we have shown the various energy components calculated using all-atom forcefield function CHARMM(Brooks et al., 2009) leads to 30% recovery of the thermostable mutants for several class A GPCRs in the top 50 predicted mutants (Fig. 3A). The CHARMM energy function includes the valence bond energy, the angle energy and the dihedral angle energy. It also includes non-bond energy terms such as Coulombic energy and van der Waals energy. In this study, we have also included other energy terms derived from statistical analysis of protein structures in the protein data bank, such as energy term favoring preferred Ramachandran backbone dihedral angles, and statistics-based term favoring salt bridges available in the ROSETTA forcefield(Alford et al., 2015). We also included a feature that classifies an amino acid as hydrophobic or hydrophilic.

### Structures of GPCRs used to calculate the features

We used the crystal structures for all the GPCRs used to train the machine learning models. For the blind test case C5aR, we generated a homology model since this work was done prior to the report of crystal structure of C5aR(Robertson et al., 2018). C5aR showed low sequence homology with existing crystal structures at the time of this modeling effort. Crystal structures of the inactive state of μ, κ, δ-opioid receptors, nociception receptor, chemokine receptors CCR5(Tan et al., 2013), CXCR1(Vaidehi et al., 2006) and CXCR4(Qin et al., 2015) showed 17 to 23% sequence homology to human C5aR. Therefore we used GPCR-I-TASSER(Zhang et al., 2015) [method to build the homology models with multiple templates. Specifically, the human C5aR sequence was first threaded through GPCR structure library to identify putative structures of fragment templates. Following which the fragments are assembled into full-length models by replica-exchange Monte Carlo simulations, which are assisted by a GPCR and membrane specific force field and spatial restraints collected from mutagenesis experiments. All models were further refined by fragment-guided dynamics simulation to eliminate inter-TM steric clashes that improve the model quality. This led to C5aR structural models with high confidence score (C-score = 2.3), especially for TM1-7 and ECL1-3 regions allowing for further use in thermostability mutation predictions.

### Structural Properties Related to Stability

Apart from energy terms, we. have also calculated structural properties of GPCRs using GPCR crystal structures where available or homology models. The interatomic structural properties we used were calculated using the Protein Contact Atlas **a**vailable at https://www.mrc-lmb.cam.ac.uk/rajini/index.html. The properties we used are (i) the betweenness and closeness centrality of each residue calculated from the crystal structures or homology models, that reflect how connected each residue is to other residues in the protein structure, (ii) the solvated area of each residue and, (iii) the property known as degree which reflects the number of residue contacts made by each residue.

### Sequence based Properties Related to Stability

To provide a comprehensive set of features to train the machine learning models, we included amino acid sequence-based property known as evolutionary coupling score(Hopf et al., 2014) that quantifies the extent to which each residue is conserved or undergoes correlated mutation. The higher the evolutionary coupling score stronger is the role of the residue in preserving the structure or function of the GPCR. To calculate the evolutionary coupling score for each amino acid in each GPCR structure we used a web based structure analysis software (http://evfold.org/evfold-web/newmarkec.do?). The evolutionary score is calculated using a multiple sequence alignment which we generated using 300 non-olfactory class A GPCR sequences and GPCRDB(Isberg et al., 2015) web server.

### Properties that discriminate the experimental data on thermostable mutants

We performed Principal Component Analysis(PCA)(Jolliffe, 2013; Tharwat, 2016) and Linear Discriminant Analysis(LDA)(Mika et al., 2003; Tharwat et al., 2017) on the 26 features listed in Table S2. These two methods reduce the multi-dimensional space into fewer dimensions. The vectors obtained from these analyses best separate the thermostable from the non-thermostable mutant data. Therefore, we have considered these vectors as additional features in training the machine learning models.

### Methods used to calculate energy related features

For the GPCRs used in the training set shown in ensembles E1 to E5 in Table S1, there were crystal structures available. However, at the time we started this work we did not have a crystal structure for the blind test case of C5aR and hence we have not used the C5aR crystal structure that has been published since(Robertson et al., 2018). Starting from the respective crystal structure for each GPCR (see Table S1 for a list of crystal structures and their respective protein databank identities) we used the LiticonDesign method(Bhattacharya et al., 2014; Bhattacharya and Vaidehi, 2012; Vaidehi et al., 2016) to generate a small ensemble of conformations for each GPCR. We have described the LiticonDesign method in detail in our previous work(Balaraman et al., 2010; Bhattacharya et al., 2014, 2008; Bhattacharya and Vaidehi, 2012). Briefly, the LiticonDesign method for predicting thermostable GPCR mutants involves two steps: (i) using a starting structural model of the GPCR, the method generates a small ensemble of conformations that allows for the perturbations in the GPCR conformation caused by mutations and, (ii) an all-atom energy function to calculate the stability of the conformations that takes into account the difference in the structural stability of the mutants and the wild type to score the positive thermostable mutants. Starting from an initial receptor structure, or homology model in the case of C5aR, all of the seven transmembrane (TM) helices are rotated simultaneously about their respective helical axes between ±5° in 10° increment. Thus, 2^7^=128 conformations are generated for the wild type receptor. Then we perform mutation of each residue to alanine (alanine in the wild type is mutated to leucine) in each of the 128 conformations and repack the side chain conformations of all of the residues using SCWRL4.0(Krivov et al., 2009), followed by steepest descent (SD) energy minimization using CHARMM27 force field for 1000 steps(Brooks et al., 2009). Each of the energy components listed in Table S2 was calculated for the wild type and the mutant for each of the 128 conformations, and averaged over the 128 conformations for the wild type and the mutant separately. Each energy component of the overall stability score was calculated as the difference in the average mutant energy of that particular energy component to the average wild type energy. For example, the van der Waals energy component of the stability was calculated as Stability score (vdw) = <E_vdw, mutant_> – <E_vdw, wild type_ > for a given single mutant where the respective energies are averaged over the 128 conformations. To give equal weight to every amino acid type in our scoring, we calculated the z score from the stability score, for every amino acid type. We will refer to this z score as the stability score from here afterwards in the paper. The stability score for each energy component and for mutant was also calculated using the Rosetta forcefield mpframework_fa_2007 as implemented in the Rosetta software(Alford et al., 2015). The different energy components listed in Table S2 were all calculated for each mutant.

### Methods used to calculate sequence and structure related features

The evolutionary coupling score is a measure of the extent of correlated evolutionary sequence changes across nonolfactory class A GPCRs and it identifies the residues that play a critical role in structural stability or function of the GPCR. We calculated the evolutionary coupling score using the EVcoupling web server(Hopf et al., 2014). The web server generated the cumulative evolutionary coupling scores for each residue in a given GPCR sequence using a multiple sequence alignment that we had provided for 300 non-olfactory class A GPCRs. We converted the evolutionary scores calculated using the webserver to z scores. The multiple sequence alignment containing a total 300 non-olfactory class A GPCRs was generated using GPCRDB(Isberg et al., 2015).

## Results

### Thermostable mutants cluster on the intracellular region of transmembrane helices 5 and 6

Four GPCRs have been subjected to comprehensive alanine scanning mutagenesis by Tate and coworkers, and the resulting mutants tested for their thermostability using a radioligand binding assay(Tate, 2012). The turkey β_1_-adrenoceptor (β_1_AR)(Serrano-Vega et al., 2008), human adenosine A_2_A receptor (A_2_AR)(Magnani et al., 2008) and human angiotensin receptor (AT_1_R) were stabilized in an inactive state that bind antagonists and with similar affinity to the wild type receptors. The rat neurotensin receptor (NTSR1)(Shibata et al., 2009) and A_2_AR(Lebon et al., 2011) were also thermostabilized in an active-intermediate state (Fig. 1A) that binds agonists with similar affinity to the wild type receptor. A total of 1231 alanine/leucine mutants were tested for thermostability out of which 256 mutants are thermostable. The criteria for thermostability we use here is if the mutant has the same or increased thermostability as the wild type receptor. The mutants with >130% of the wild type are *strong* thermostable mutants. Analysis of the positions of thermostable mutations, show that they are distributed throughout the TM helices and loop regions with a higher percentage located in TM5, TM6 and intracellular loop 3 (ICL3) (Fig. 1B and Fig. S1A). The thermostabilizing mutations common to β_1_AR, both conformational states of A_2_AR and NTSR1, are clustered in the intracellular region of TM5, TM6 and ICL3 (Fig. 1C and Fig. S1B). The highly conserved residue in each TM helix is numbered as the 50^th^ residue according to Ballesteros-Weinstein residue numbering scheme for GPCRs(Ballesteros and Weinstein, 1995). The clustering of mutations in this mobile domain of the GPCR highlights that GPCRs have evolved to retain flexibility in their dynamic equilibrium, and mutation away from natural sequence effectively shunts this conformational flexibility. Understandably, mutation of this most conserved residue in each TM helix decreases the stability of the receptor (Fig. S2). This may be related to the mechanism of receptor activation, which involves significant changes in the orientation of TM5 and TM6 with respect to TM3 (Fig. 1A). However, mutation of other conserved residues such as Asn or Tyr of the NPxxY motif on TM7 does not always destabilize the receptor (Fig. S2B). GPCRs are in a dynamic equilibrium between inactive and active states, so thermostabilization of an inactive state may arise through mutations that prevent the transition to active states or destabilize the active states. Other mutations may stabilize the receptor through increasing contacts between residues, promoting formation of inter-helical backbone hydrogen bonds and/or entropic effects(Lee et al., 2015; Vaidehi et al., 2016). Mutation of aliphatic hydrocarbon amino acids such as Val, Ile, Leu, Met and Cys more often lead to thermostabilization than other types of amino acids (Fig. 1D).

### A Machine Learning Model for predicting GPCR thermostabilizing mutations

Comparison of the amino acid positions of thermostabilizing mutations in different receptors shows that there is no significant conservation between amino acid position and thermostability. Thus, transferring thermostabilizing mutations between GPCRs works only if the receptor sequences are highly conserved, for example, between the turkey β_1_AR and human β_1_AR(Serrano-Vega and Tate, 2009). Therefore, a robust mutagenesis prediction method will help scientists reduce time and cost, focusing the number of trials on more promising amino acids mutations. Using the largest set of experimental thermostability data publicly available, we have employed ensemble ML models for predicting thermostabilizing mutations ahead of experiments. We developed a workflow as shown in Fig. 2 to train and optimize ML models. The two major sections of this workflow consist of: (i) preparing the experimental data for machine learning (ii) calculating features describing thermostability of GPCRs. Detailed description of these two sections are given in the Methods section.

**Figure 2:**
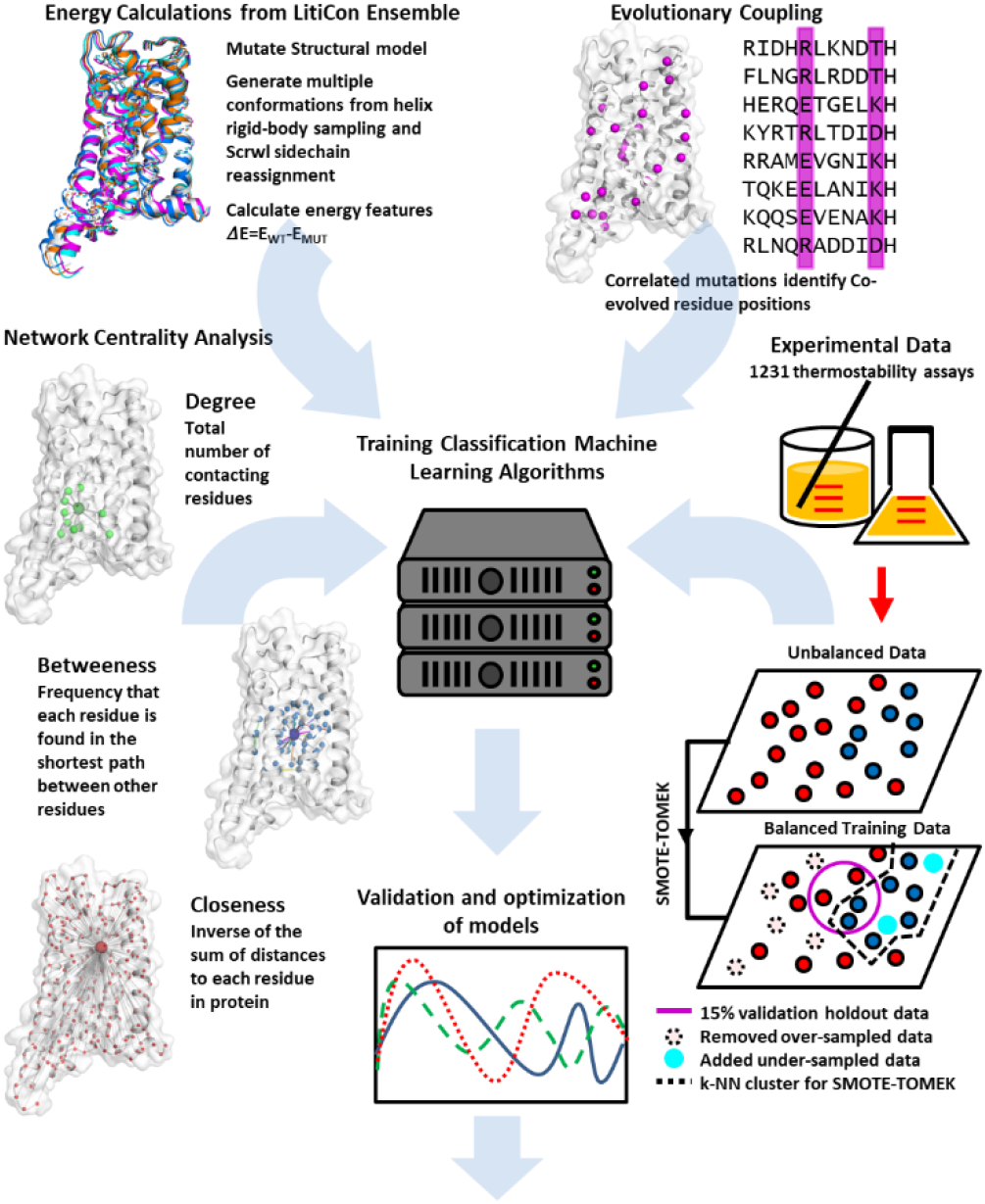
Workflow developed for optimizing, validating and testing the machine learning methods. The figure shows the atomic level features that we have calculated to recapitulate the GPCR structural thermostability. Properties based on co-variation of amino acid sequence positions, structural properties such as betweeneness, centrality calculated from network analysis, and thermodynamic energies calculated from structural ensemble. The right side of the figure shows the algorithm we used for balancing the experimental thermostability that was the target data for training the machine learning models.

### i. Preparing experimental data for ML optimization

We divided the five sets of experimental thermostability data set on four GPCRs (A_2A_R has two sets of data, one for the antagonist bound inactive state and other for the agonist bound active-intermediate state (Fig. 1A)) into five ensembles shown in Table S1. In each of the five ensembles we trained the ML models using four datasets and held out the fifth dataset as a test for prediction. The experimental thermostability data on 1231 mutants contains 256 thermostable mutants and 975 non-thermostable mutants and hence is not balanced. Therefore, to balance the training data we used the Synthetic Minority Oversampling Technique (SMOTE)-Tomek sampling(Batista et al., 2004) method to both under sample the majority class (non-thermostable mutants) and oversample the minority class (thermostable mutants) simultaneously. SMOTE oversamples the minority class using its k nearest neighbors(Guo et al., 2010) (KNN). Tomek removes samples from the majority class by measuring intrasample distance and removing outliers. 85% of the resulting balanced data was used for training the ML models, while 15% data was held out from training, for ML model validation.

### ii. Features derived from sequence, structure and thermodynamics describe the thermostability of GPCRs

Features are individual, measurable properties that describe the characteristics of the phenomenon being observed, thermostability of mutants in this case. Identifying informative, discriminating, and independent GPCR features that best describe its thermostability is a crucial step for effective ML classification algorithms. As listed in Table S2, we have used features extracted from class A GPCR amino acid sequence analysis, structural analysis and atomistic energy calculated from conformation ensembles.

### Sequence and Structural features that describe the thermostability

We performed evolutionary covariation analysis using the “EV couplings” webserver (EVfold.org)(Hopf et al., 2014) with the multiple amino acid sequence alignment of 300 non-olfactory class A GPCRs as input, generated using the toolkit from “GPCRdb”(Isberg et al., 2015). This analysis calculates the covariation of any pair of amino acid positions in the class A GPCR sequence alignment. We have abstracted structural properties using a graphical network model of the GPCR crystal structures where available or homology models if the crystal structures are not available. We used the “Protein Contact Atlas” web server (https://www.mrc-lmb.cam.ac.uk/rajini/index.html) and calculated residue based structural properties such as degree, centrality, betweenness, and closeness, from a single input of the structure or structural model of the GPCRs. The solvent accessible surface area (referred to as solvated area) for each residue was calculated using the POPS(Cavallo et al., 2003) algorithm within “Protein Contact Atlas” (https://www.mrc-lmb.cam.ac.uk/rajini/index.html). More details are given in the Methods section. Together, these structural properties capture the non-covalent interactions between residues in the protein framework.

### Energy features calculated from forcefield function

In our previous studies we used the LiticonDesign(Bhattacharya et al., 2014; Bhattacharya and Vaidehi, 2012; Vaidehi et al., 2016) method to generate an ensemble of conformations for each GPCR using the rotational degrees of freedom of the seven TM helices. The stability score was calculated as the difference between the average value of the energy component of the mutant (averaged over the ensemble) and the average energy component for the wild type. For example, Stability score = <E_vdw, mutant_> – <E_vdw, wild type_ > for the van der Waals component. The energy components were calculated using atomic forcefield functions, CHARMM27(Brooks et al., 2009) and ROSETTA(Alford et al., 2015), to describe the thermostability of GPCRs. Previously, we had shown that using an ensemble of conformations for the stability score lead to better recovery of known mutants compared to using a single structural model(Bhattacharya et al., 2014).

### Merits of the features in recovering thermostable mutants

We assessed the merit of each of these features to predict thermostable mutants by using the z-scores of each feature to test how well they recover the thermostable mutations in the 1231 mutant dataset. The number of thermostable mutations recovered in the top 50 predicted mutants using each of the sequence, structural and energy-based features as well as their sum are shown in a radial plot (Fig. 3A). All the features recover more thermostable mutants in the top 50 predicted mutants compared to the random predictions, demonstrating the merit of these properties in describing the GPCR thermostability. The sum of the z-scores of all the 26 features recovers nearly three-fold the number of thermostable mutants compared to the individual features since the individual features recover distinct set of thermostable mutants. This suggests that the sum of the properties captures the GPCR thermostability better than a single property.

**Figure 3:**
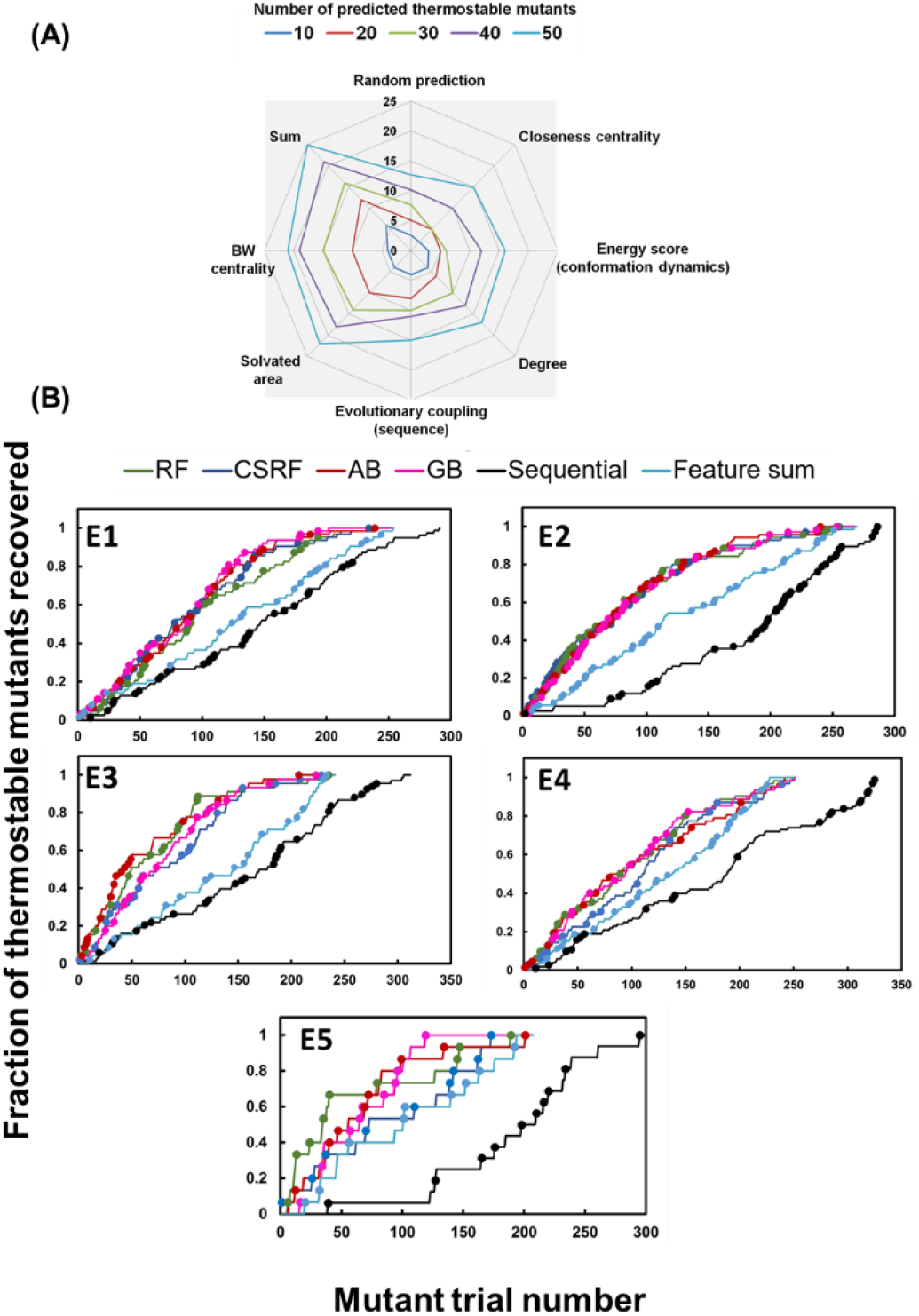
**A**. Merits of the sequence, structure and dynamical based properties used for training the ML models. The recovery of thermostable mutants greater than 100% of the wild type stability using each of the features. **B**. The five figures here show the fraction of thermostable mutants recovered using the predicted prioritized listof mutants using different machine learning models on the five ensemble of data used for training and testing the four machine learning models in this work (for breakdown of the GPCR systems included in training and testing see Table S1). The aqua colored curve in this figure is the recovery rate using the sum of the features without any machine learning.

### Machine Learning Algorithms Tested

Our goal is to use machine learning methods to recover the maximum number of thermostable mutants in fewer mutant trials. We used supervised “classification” learning models to identify the relationship between our feature set and the thermostability class. We labeled our training data of experimental mutation results as either class “1” (thermostable) or class “0” (non-thermostable) when compared to the wild type. We have tested the performance of four machine learning algorithms: random forest(Breiman, 2001) (RF), cost sensitive random forest(Elkan, 2001) (CSRF), adaptive boosting(Freund and Schapire, 1999) (aka AdaBoost) (AB), and gradient boost(Hastie et al., 2009) (GB) on 5 ensembles. To achieve the optimal parameters that will allow the model to generalize to new data, we performed hyper-parameterization using the 85% of the balanced data for training and the 15% for validation. We used the Matthews Correlation coefficient (MCC)(Boughorbel et al., 2017) as a metric to assess the performance of each of the ML models since it is a composite function of the true positives (TP), true negatives(TN) and false positives(FP) and false negatives (FN). The performance of the four ML models was validated on the 15% held out data from the balanced training data. We further tested the hyper-parametrized ML models on the test protein that was not used in the training, in each of the five ensembles (Table S1). Fig. S3A and B show the MCC ratios for the validation data and for the test proteins respectively. The Random Forest and AdaBoost models perform better than the Cost Sensitive Random Forest and Gradient Boost models in most cases with a higher MCC value. These two models predict more true positives and negatives, while having a lower prediction rate of false positive and negatives. Figure 3B shows the fraction of thermostable mutations retrieved as a function of the prioritized ML-predicted alanine mutation list as compared to the systematic alanine scanning mutation list. We observe that 35% to 40% of the thermostable mutants are recovered in the top 50 prioritized mutant list compared to less than 10% in the systematic alanine scanning mutations. We note that the systematic alanine scanning mutations list contains only those residues for which we have reliable experimental thermostability results.

### Blind predictions of thermostable mutations for C5aR

Using the four optimized ML models, we performed blind predictions on the thermostabilizing single point mutants for the inactive state of wild type C5aR GPCR. We predicted the thermostable mutants using the four ML models not previously trained or validated on C5aR. Simultaneously, systematic alanine mutagenesis experiments were performed to identify thermostable mutants for the antagonist-bound inactive state of C5aR using the procedure described in Methods section.

We generated homology based three-dimensional structural models for C5aR, followed by the LiticonDesign method(Balaraman et al., 2010; Bhattacharya et al., 2014; Bhattacharya and Vaidehi, 2012) to generate a small conformation ensemble starting from the homology model. We calculated the energy features listed in Table S2, for all the alanine scanning mutations using ROSETTA(Alford et al., 2015) and CHARMM 27(Brooks et al., 2009) forcefields. We calculated the evolutionary coupling scores for C5aR using non-olfactory class A GPCRs multiple sequence alignment, and the structural features using the homology model. These features were used in the ML models to calculate the probability score for mutating each residue in C5aR to alanine (or mutate to leucine if the wild type is alanine). We compared our blind predictions to the experimental thermostabilities that we obtained from our collaborators at Pfizer. Fig. 4A shows the fraction of thermostable mutants recovered from the ala scanning mutation list prioritized by each ML model (green, blue, red, purple) compared to performing the sequential alanine scanning mutations in C5aR (black). The dots plotted in Fig. 4A are the positions at which strong thermostable mutants (>130% wild type ligand-binding density) are recovered. All four ML models recover the thermostable mutants at a higher recovery rate than the systematic alanine scanning mutation trials, but Random Forest and Adaptive Boost models recover consistently higher number of thermostable mutations, even for as few as 25 mutation trials (Fig. 4B). The gradient boost model recovers 14 thermostable mutants containing 4 strong thermostable mutants in the top 50 prioritized list. Thus, we demonstrate that sequence, structure, and dynamics derived features are useful tools for efficiently prioritizing thermostabilizing mutations.

**Figure 4:**
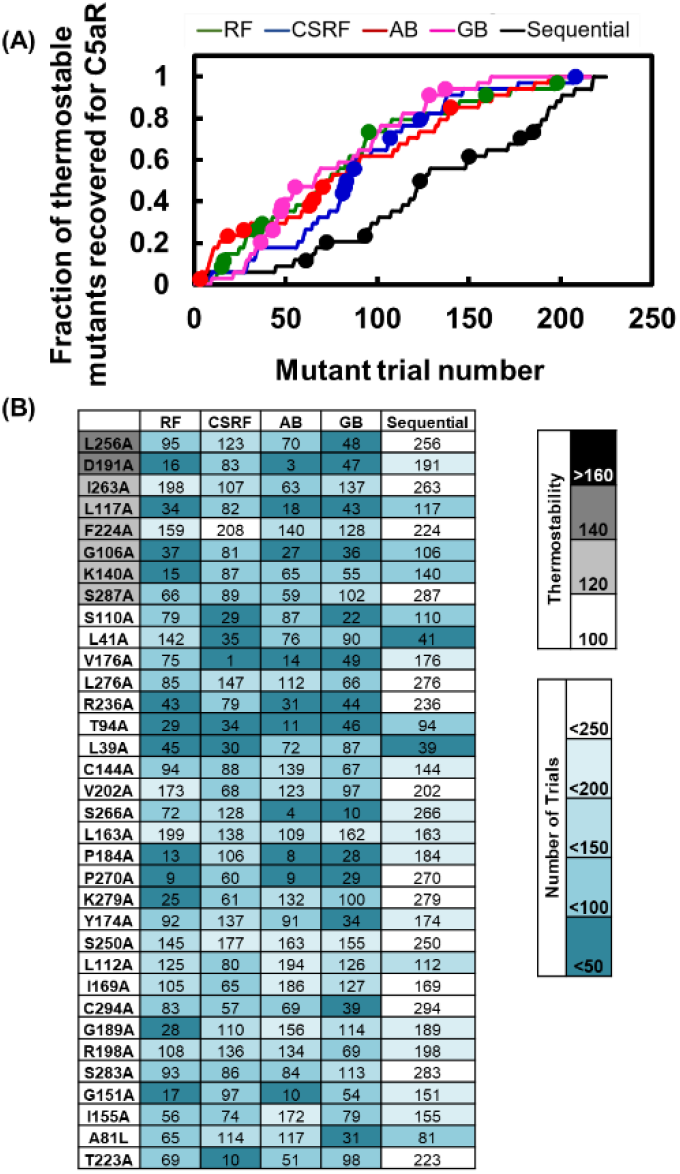
**A**. Recovery rate of thermostable mutants from the prioritized list of alanine scanning mutations for the blind test on C5aR, prioritized using different machine learning models. The x axis shows a prioritized list of mutations predicted by the four machine learning models. The black curve is the recovery rate when performing systematic alanine scanning mutations along the sequence. **B**. This table shows the mutant trial numbers at which the true positive thermostable mutations were recovered. The strength of thermostability are shown in grey scale. The sequential trial numbers shown in the last column of this table assumes that the mutation trial will start from amino acid 1 in the sequence,

### Prediction of thermostable mutants for agonist bound active-intermediate state of the adenosine A_2A_R

Magnani et al(Magnani et al., 2008) identified single point thermostable mutants of the agonist-bound (NECA) active-intermediate state and the antagonist-bound (ZM-241385) inactive state of A_2A_R using alanine scanning mutagenesis across the entire receptor sequence. They identified mutations that retained the ligand binding at higher temperatures as compared to the wild type(Magnani et al., 2008). The majority of thermostabilizing mutations are distinct to either the inactive or active-intermediate state of the receptors, with few common mutation positions (Fig. S4). Mutants present in transmembrane helices TM1 to TM4 (shown in magenta, Fig. S4), stabilize the antagonist-bound inactive state, while mutants that specifically stabilize the agonist-bound active intermediate state are predominantly found in TM5, TM6, and TM7 (shown in yellow, Fig. S4). This shows that the thermostable mutants are not easily generalizable, even for the same receptor in different conformational states.

We tested the optimized ML models in predicting thermostabilizing mutations for the agonist bound active-intermediate state of A_2A_R. We trained the ML models on the experimental thermostabilizing data for inactive state of β_1_AR, inactive state of A_2A_R, inactive state of AT1R, and agonist-bound active-intermediate state of NTSR1. Fig. 5A shows the recovery curve for agonist-bound active-intermediate state of A_2A_R using the ML models trained on the ensemble of thermostability data. The ML models recover 17 to 20 thermostable mutants out of which 11 to 14 are strong thermostable mutants in the top 50 prioritized alanine mutants as shown in Fig. 5B. This test validates that our ML prioritized list recovers thermostable mutants much faster than systematic alanine scanning.

**Figure 5:**
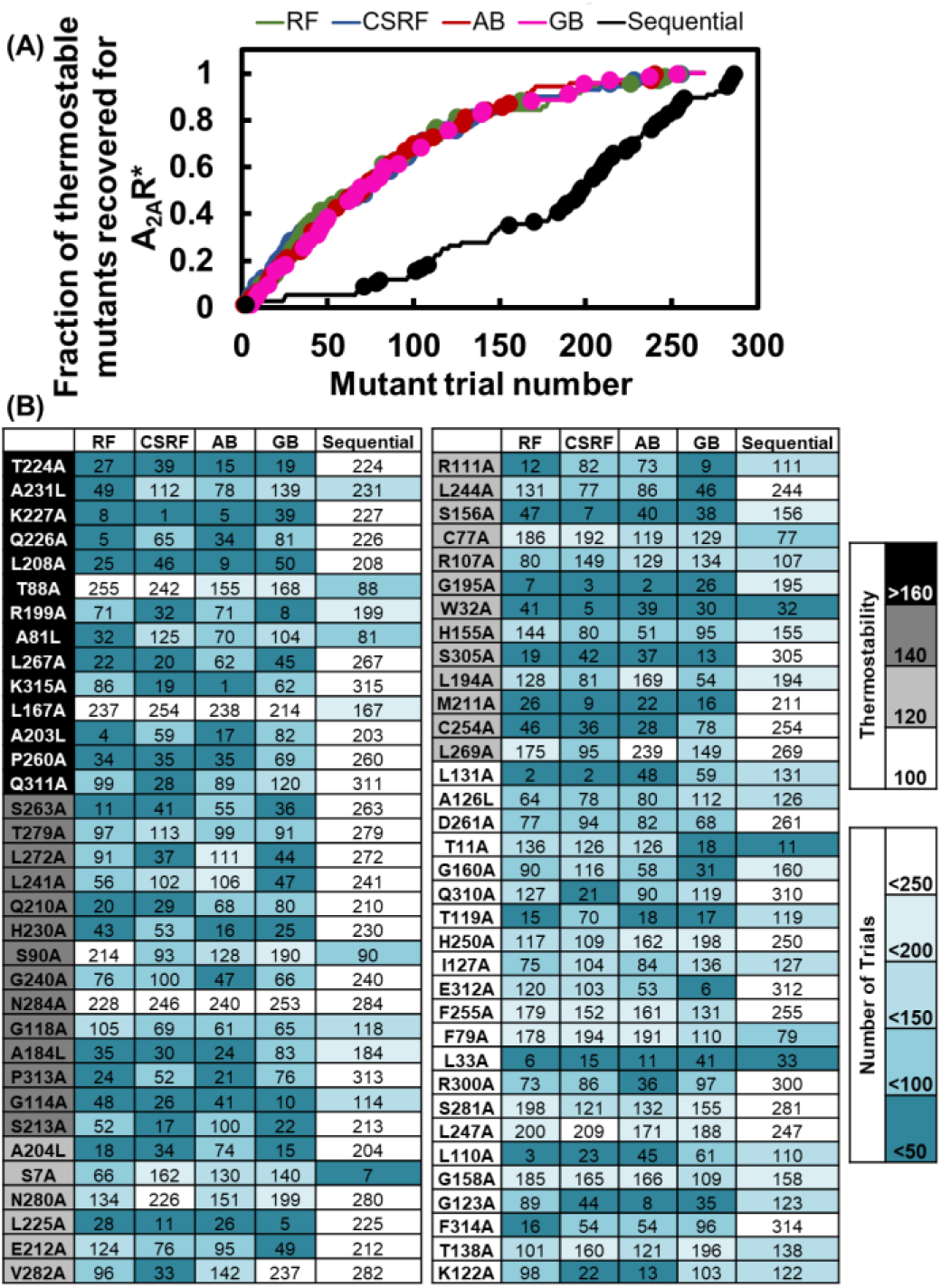
**A**. Recovery rate of thermostable mutants from the prioritized list of alanine scanning mutations for the agonist bound active-intermediate state of A_2A_R, prioritized using different machine learning models. The black curve is the recovery rate when performing systematic alanine scanning mutations along the sequence. **B**. This table shows the mutant trial numbers at which the true positive thermostable mutations were recovered. The strength of thermostability are shown in grey scale. The sequential trial numbers shown in the last column of this table assumes that the mutation trial will start from amino acid 1 in the sequence,

### Relative importance of features describing thermostability

The most important features for thermostability prediction using a ML model can be determined via feature selection (model agnostic) or directly from the ML model by feature ranking. For three of the four ML models (Random Forest, AdaBoost and Gradient Boost) the feature ranking was generated using the importance measure, “Mean Decrease in Impurity (MDI)” otherwise known as the “Gini Importance” (defined in the Methods section). Across the three ML models, Fig. 6A shows that the “solvated area” feature is the most important and removal of this feature from the list, shows poor recovery of thermostable mutants (Fig. 6B).

**Figure 6:**
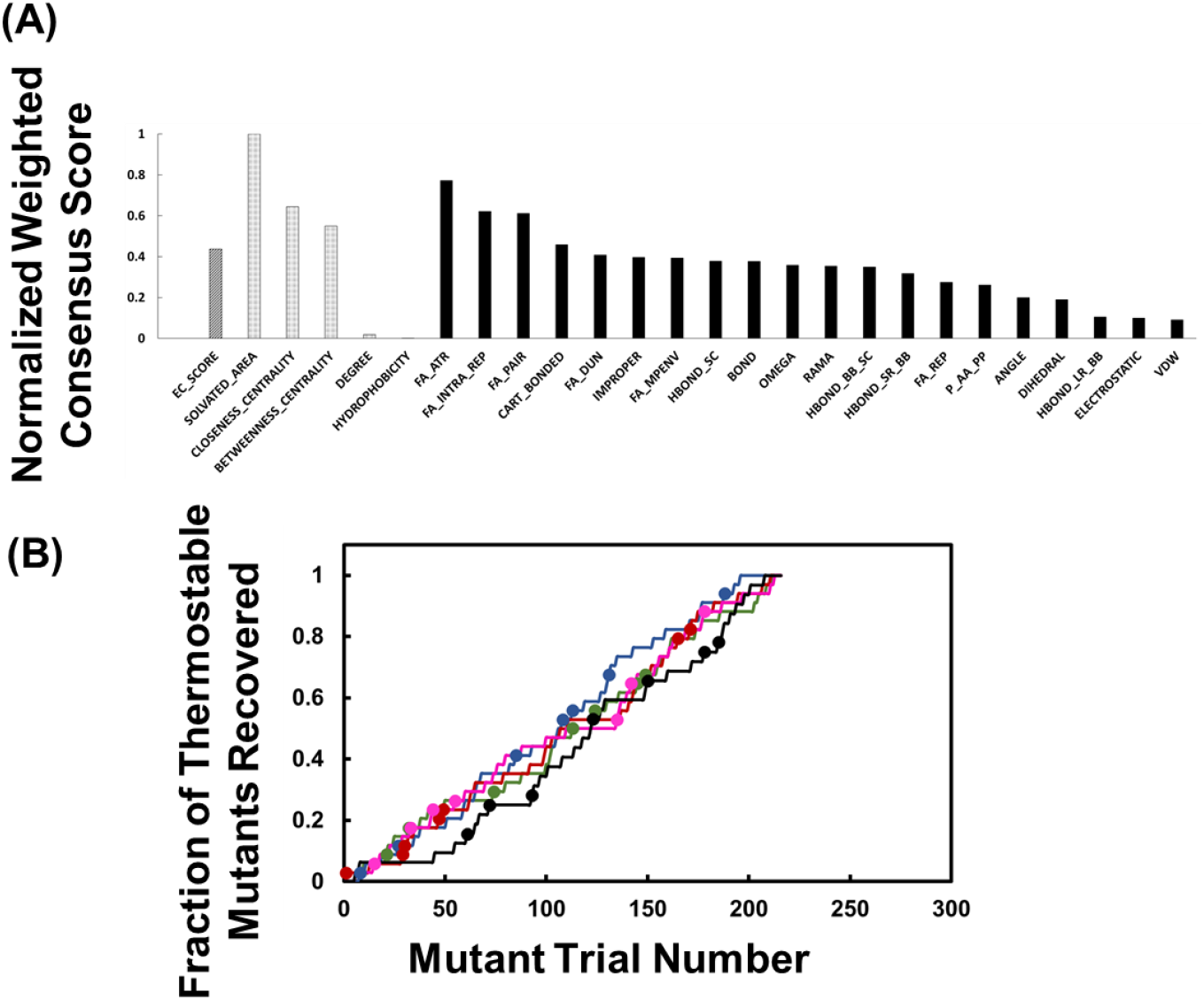
**A.** Ranking the weights of the 26 features derived from sequence, structure and thermodynamics energy analysis, used to describe the GPCR thermostability. The 26 features used here are described in Table S2 of the Supporting Information. On the y-axis is the normalized weighted consensus score derived from feature ranking for each machine learning model. The normalization was done with respect to the solvated area feature. **B.** The percentage recovery rate of thermostable mutants using 25 features after omitting the high weighted “solvated area” feature.

## Discussion

In this study, we have used an innovative combination of molecular level features calculated from GPCR sequence co-variation analysis, structural features calculated from network analysis and thermodynamic ensemble properties of GPCRs to recapitulate the thermostability of GPCRs. As shown in Fig. 3, the sum of these features, without any machine learning methods involved, recover more thermostable mutations than a random prediction or systematic alanine scanning mutation analysis along the sequence of the GPCR. These 26 features when combined with four different classification based machine learning methods provide robust models for prediction of thermostabilizing mutations as demonstrated in the blind test case of C5aR. These ML models are useful in prioritizing alanine scanning mutations for experimentalists. The trained ML models performed better than random predictions or systematic alanine scanning mutations along the amino acid sequence of C5aR. These predictions demonstrate the power of combining physically meaningful features of GPCRs with machine learning models in reducing the time and reagent costs for experiments. We have demonstrated that the features calculated from a homology model of the GPCR structures works well for the blind test case C5aR.

We have used the largest publicly available thermostability dataset (1231 mutants) for training the ML models. These data are for the inactive state of β_1_AR, inactive state of A_2A_R, inactive state of AT1R, and active-intermediate state of A_2A_R and NTSR1. As anticipated in many biological datasets, our thermostability data was imbalanced by negative data, which complicates the development of a classification model generalizable to new data(Rahman and Davis, 2013). Using SMOTE-TOMEK(Batista et al., 2004) to balance our training data resulted in a significant improvement in MCC.

### Predictions for thermostabilizing mutants of different conformation states

We have demonstrated a high recovery rate of thermostable mutants for GPCRs in different conformational states. Machine learning models trained on all data show a recovery of over 30% of thermostable mutants (and 20% of ‘highly’ thermostable mutants) within the top 50 prioritized mutations. We note that the structural and energetic features describing the stability of the agonist bound state were calculated using the structural models for active-intermediate state rather than inactive states. We infer that it is important to calculate the features using the structural model of the conformation state for which thermostabilization is desired. Particularly, we believe individual features may capture distinct aspects of the phenomenon of thermostability, but the right ensemble of features is critical for better predictive ability.

## Author Contributions

S.A. and N.V. designed the study, S.G. generated the features for describing the thermostability and tested them, S.M. and S.A. developed the workflow and tested the machine learning models, S.M., S.G., S.A., and N.V. analyzed the machine learning results S.M., S.G. and M.S. generated the figures, X.C., X.Y., M.C.G., K.F.F., Y.C., V.S. and X.Q. performed the experiments and contributed to the analysis of the blind test results for C5aR, C.G.T. shared the experimental thermostability data for 4 GPCRs, S.M., S.G., S.A., M.S., and N.V. wrote the manuscript.

## Acknowledgements

This work was funded by NIH grant R01-GM097261 to N.V.

